# Outcome of *Drosophila* microbiota manipulation depends on dietary preservative formula and batch variation in dietary yeast

**DOI:** 10.1101/2023.01.30.526386

**Authors:** David R. Sannino, Adam J. Dobson

## Abstract

Gut microbiota are fundamentally important for healthy function in their animal hosts. The fly *Drosophila melanogaster* is a powerful system for understanding the underlying host/microbe interactions, with modulation of the microbiota inducing phenotypic changes that are conserved across animal taxa. The context-dependence of these responses has not been explored systematically, which may confound repeatability. Here we show that the microbiota’s impact on fly triacylglyceride (TAG) levels - a commonly-measured metabolic index - depends on factors in fly media that are rarely considered or controlled, and are not standardized among laboratories: media preservative formula, yeast batch, and the interactive effect of their combinatorial variation. In studies of conventional, axenic and gnotobiotic flies, we found that microbial impacts were apparent only on specific yeast-by-preservative conditions, with TAG levels determined by a tripartite interaction of the three experimental factors. When comparing axenic and conventional flies, we found that preservatives rather than microbiota status was the main driver of variance in host TAG, and certain yeast-preservative combinations reversed microbiota effects on TAG levels. Further, comparisons between TAG levels of axenic flies and those associated with *Acetobacter pomorum* or *Levilactobacillus brevis* determined that preservatives, microbiota status, and their interaction were the major drivers of TAG variation. Our results suggest that the microbiota shapes the host TAG response in a manner dependent on the combination of the dietary factors of preservative formulation and yeast batch, with implications for repeatability, interpretation, and optimal experimental standards.

**Importance:** *Drosophila melanogaster* is a premier model for microbiome science, which has greatly enhanced our understanding of the basic biology of host-microbe biology. However, often overlooked factors such as dietary composition, including yeast batch variability and preservative formulation used, may cofound data interpretation of experiments within the same lab and lead to different findings when comparing between labs. Our study supports this concept; we find that host TAG levels are not solely dependent on the presence or absence of microbiota members, but rather the combinatorial effects of microbiota members, yeast batch, and preservative formulation used, with preservatives being the largest driver. It serves as a cautionary tale that underappreciated components of fly rearing can mask or drive phenotypes that are believed to be impacted by microbiota members.

## Observation

Fruitflies are a preeminent model for understanding fundamental host-microbiome biology, thanks to experimental tractability, powerful molecular tools, and a simple microbiota dominated by culturable *Lactobacillaceae* and *Acetobacteraceae* (1) (2). Flies can be routinely made germ-free (axenic), or selectively reassociated with defined cultures of physiologically- and ecologically-relevant microbiota (gnotobiotic). The fly microbiota is less complex than in vertebrates, yet effects on a plethora of host traits are conserved (3–14), potentially indicating common mechanisms that can be rapidly characterized in the fly.

The microbiota affect fly nutrition, and so variation in microbiota and diet have mutually- interdependent effects (15). Brewer’s yeast is ubiquitously included in fly diets (16). Importantly, yeast is commercially supplied in lots originating from distinct production batches, with potentially variable chemical composition. This potentially introduces nutritional inconsistencies among distinct lots (16), that may modify response to microbiota manipulation.

Fly diets also commonly contain antimicrobial preservatives. Preservative formulae vary both in composition and concentration, and in some microbiota studies they are omitted entirely (10, 13, 14, 17). The commonly-used preservative nipagin (methylparaben) affects the density of *Acetobacter* (18), which may alter growth in fly food, and thereby modify physiological impact. Also nipagin is dissolved in ethanol, which interacts with variation in the microbiota (19). Acid preservatives are also used, which may modulate fly function though effects on the microbiota (e.g. density, metabolic substrate provision), diet (e.g. pH and nutrient solubility), and direct effects on the fly (20).

Here we test whether physiological impact of altering the fly microbiota depends on dietary yeast batch and preservatives. We used two lots of one supplier’s yeast, denoted A or B. We either omitted preservatives, or added (1) phosphoric acid and propionic acid (15), or (2) nipagin and propionic acid (13). These ingredients were incorporated into otherwise identical sucrose-yeast-agar (SYA) diet (21). We measured triacylglyceride (TAG) levels, the main storage lipid, which are commonly measured as a metabolic index due to interest in the microbiome’s role in human obesity (22).

First, we applied a simple microbiome manipulation, comparing conventionally-reared and axenic females, three days after adult emergence. We analyzed data with ANOVA (Table 1) and *post-hoc* tests (Table 2). TAG response to bacterial elimination depended on the interaction of yeast batch and preservative formula (ANOVA: bacteria*yeast*preservative F_2,106_=3.73, p=0.03; Table 1). This interaction obscured the anticipated increase of TAG in axenics (ANOVA: bacteria F_1,106_=0.54, p=0.46, Table 1), suggesting that microbial capacity to modulate TAG depends on a yeast*preservative interaction. To examine specifically how, we stratified our analysis per yeast*preservative combination. Without preservatives, on both yeasts, TAG was elevated in axenics (Table 2). Surprisingly, this response was reversed by a specific yeast*preservative combination, with conventionals showing higher TAG than axenics on yeast A and with preservative formula 2 (Table 2, Figure 1A). Furthermore, microbial manipulation did not affect TAG in any other condition including preservatives, on either yeast (Table 2). Interestingly, preservative formula 2 increased TAG even in axenic flies, but only on yeast B (Table 2), suggesting effects via fly or food. Further, the TAG levels were typically more variable when preservatives were present on both yeasts, and this variability was most pronounced on yeast B with preservative set 2 (Figure 1A). To determine relative contributions of experimental factors, we calculated effect sizes (Eta^2^) for each experimental variable and their interactions (Figure 1B). This indicated that preservatives were the biggest source of variance (Figure 1B). Confidence intervals for all other terms overlapped, suggesting equivalent contributions to overall variation. These results indicated that preservatives, and their interaction with yeast batch, are a hitherto unappreciated factor determining fly TAG, which can eclipse effects of microbiota.

**Figure 1.**
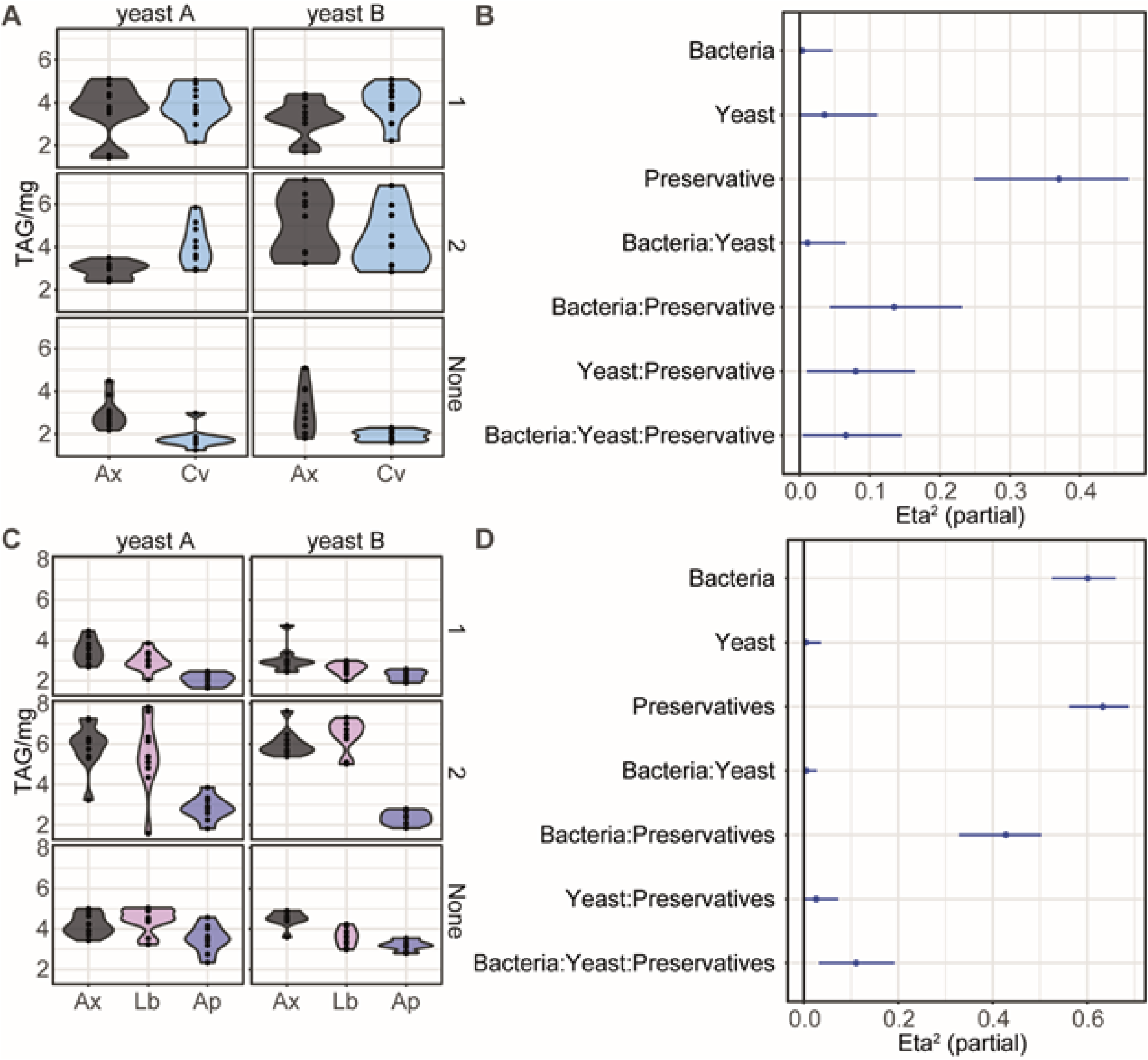
Metabolic impact of microbiota depends on combination of yeast batch and preservative formula. **(A)** and **(C)** are panel violin plots comparing TAG levels, separated by yeast batch A (left-side) or B (right-side) and by preservative formulation: 1 (top), 2 (middle), or none (bottom). **(A)** Comparisons between axenic (Ax) and conventional (Cv) flies show that TAG levels are only reduced in conventional flies when no preservatives are present. **(C)** Comparisons of TAG levels between axenic (Ax), *Levilactobacillus brevis* DmCS003 (Lb), and *Acetobacter pormoum* DmCS004 (Ap) associated flies show that Acetobacter pomorum reduces TAG levels relative to axenic flies in most conditions, with there being no difference between axenic and L. brevis associated flies. Preservative set 2 leads to the greatest difference in TAG levels as axenic and L. brevis flies have elevated TAG levels in comparison to their counterparts raised on food lacking preservatives or containing set 1. (B) and (D) Effect size (Eta2) plots looking at the contribution bacteria, yeast, preservatives, and their interactions have on accounting for variance. **(B)** In experiment 1 comparing axenic and conventional flies, preservatives are the major contributor of variance as all other factors have overlapping confidence intervals. **(D)** In experiment 2 comparing axenic to monoassociated flies, bacteria and preservatives are equally major contributors to the variance in TAG observed, with their interaction being another significant contributor.

**Table 1.** ANOVA (type 3) testing for preservative*bacteria interactions that determine lipidemia (TAG μg mg^-1^) in conventionally-reared versus axenic flies.

| Term | Sum Sq | Df | F | Pr(>F) |
| --- | --- | --- | --- | --- |
| (Intercept) | 1357.49 | 1 | 1484.85 | <2.20E-16 |
| Bacteria | 0.49 | 1 | 0.54 | 0.46 |
| Yeast | 4.18 | 1 | 4.57 | 0.034 |
| Preservative | 56.17 | 2 | 30.72 | 2.90E-11 |
| Bacteria:Yeast | 1.17 | 1 | 1.28 | 0.26 |
| Bacteria:Preservative | 15.35 | 2 | 8.40 | 0.0004 |
| Yeast:Preservative | 8.64 | 2 | 4.73 | 0.011 |
| Bacteria:Yeast:Preservative | 6.83 | 2 | 3.73 | 0.027 |
| Residuals | 96.91 | 106 |  |  |

**Table 2.** Effects of microbiota (conventional vs. axenic) on TAG levels of flies reared on specific combinations of dietary yeast and preservatives.

| Yeast | Preservative | estimate | SE | df | t ratio | p |
| --- | --- | --- | --- | --- | --- | --- |
| A | 1 | -0.231 | 0.428 | 106 | -0.541 | 0.5898 |
| B | 1 | -0.748 | 0.428 | 106 | -1.748 | 0.0833 |
| A | 2 | -1.131 | 0.439 | 106 | -2.575 | 0.0114 |
| B | 2 | 0.599 | 0.428 | 106 | 1.401 | 0.1642 |
| A | None | 1.152 | 0.439 | 106 | 2.622 | 0.0100 |
| B | None | 1.135 | 0.428 | 106 | 2.653 | 0.0092 |

*Acetobacter* and *Lactobacilliaceae* have strain-specific effects on fly physiology (10): Monoassociation with *Acetobacter spp*., but not *Lactobacilliaceae*, recapitulates conventional fly TAG levels (10), and growth of *Acetobacter* but not *Lactobacilliaceae* is impacted by nipagin (18). Could yeast*preservative*microbiota effects on the fly be driven by particular bacterial strains? We made gnotobiotic flies with *A. pomorum* (DmCS004) and *L. brevis* (DmCS003), and axenic controls, and modulated yeast and preservatives, to determine strain*yeast*preservative effects (Figure 1C), and again applied ANOVA (Table 3) and *post-hoc* analyses (Tables 4–5). TAG response to bacterial elimination again depended on the interaction of yeast batch and preservative formula (ANOVA: bacteria*yeast*preservative F_4,162_=5.03, p=0.0008; Table 3). Across all preservative and yeast conditions, *A. pomorum* gnotobiotes had lower average TAG than axenics and *L. brevis* gnotobiotes (Figure 1C). Preservative set 2 visibly elevated TAG levels in axenic and *L. brevis*-associated flies (Figure 1C). To assess strain-specific impacts of yeast*preservative, we stratified our ANOVA analysis by bacteria (Table 4), revealing yeast*preservative effects in gnotobiotes with *L. brevis* (F_2,162_=9.411, p=0.0001), but not with *A. pomorum* (F_2,162_=1.180, p=0.3098) or in axenic flies (F_2,162_=1.653, p=0.1946). Preservative variation had a significant effect across all microbial conditions (Table 4), while yeast had no significant effect in any microbial condition (Table 4). We also examined among-strain differences, stratified by yeast*preservative (Table 5). We expected that *Acetobacter* sensitivity to preservative set 2 (containing nipagin (18)) would impair rescue of TAG by *A. pomorum*, but, on the contrary, *A. pomorum* still rescued TAG when nipagin was present, on both yeasts (Table 5). In fact our measure of difference (t ratio) was consistently highest when preservatives contained nipagin (set 2). Thus, *A. pomorum* nipagin sensitivity did not appear to translate into impaired modulation of host TAG. Eta^2^ (effect size) calculations indicated that preservative formula and bacterial strain were the leading contributors to TAG variation in this experiment (Figure 1D). The preservative*bacterial strain interaction had a substantially-sized (and statistically significant: p<2.2e-16, Table 3) effect, suggesting that variation in bacterial strain and preservatives conspired to produce sizeable variation. Altogether, these results indicated that (1) impacts of varying microbiota strains depend on yeast*preservative variation, (2) the lower- order preservative*bacterial strain interactions was a particularly large source of variation, and (3) the effect of changing preservatives is equivalent to the effect of perturbing the microbiota.

**Table 3.** ANOVA (type 3) testing for preservative*bacteria interactions that determine lipidemia (TAG μg mg^-1^) in flies reared either axenically, or in association with *A. pomorum* or *L. brevis*.

| Term | Sum Sq | Df | F | Pr(>F) |
| --- | --- | --- | --- | --- |
| (Intercept) | 2633.33 | 1 | 5516.35 | <2.20E-16 |
| Bacteria | 116.37 | 2 | 121.89 | <2.20E-16 |
| Yeast | 0.33 | 1 | 0.68 | 0.41 |
| Preservatives | 133.4 | 2 | 139.73 | <2.20E-16 |
| Bacteria:Yeast | 0.38 | 2 | 0.40 | 0.67 |
| Bacteria:Preservatives | 57.81 | 4 | 30.27 | <2.20E-16 |
| Yeast:Preservatives | 2.09 | 2 | 2.19 | 0.12 |
| Bacteria:Yeast:Preservatives | 9.6 | 4 | 5.03 | 0.0008 |
| Residuals | 77.33 | 162 |  |  |

**Table 4.** Impact of yeast*preservative interactions under specific microbiota conditions. Joint tests are applied, stratified by microbial association.

| Bacteria† | term | df1 | df2 | F.ratio | p.value |
| --- | --- | --- | --- | --- | --- |
| Ax | Yeast | 1 | 162 | 0.014 | 0.9047 |
| Ax | Preservatives | 2 | 162 | 77.937 | <.0001 |
| Ax | Yeast:Preservatives | 2 | 162 | 1.653 | 0.1946 |
| Lb | Yeast | 1 | 162 | 0.167 | 0.6833 |
| Lb | Preservatives | 2 | 162 | 106.995 | <.0001 |
| Lb | Yeast:Preservatives | 2 | 162 | 9.411 | 0.0001 |
| Ap | Yeast | 1 | 162 | 1.308 | 0.2545 |
| Ap | Preservatives | 2 | 162 | 15.347 | <.0001 |
| Ap | Yeast:Preservatives | 2 | 162 | 1.18 | 0.3098 |

† Ax = axenic, Ap = *Acetobacter pomorum* DmCS004, Lb = *Laevilactobacillus brevis* DmCS003

**Table 5.** Differences in TAG levels among gnotobiotic and axenic flies reared on specific combinations of dietary yeast and preservatives.

| Yeast | Preservative | Contrast<br>(bacterial<br>association)<sup>†</sup> | Estimate | SE | df | t ratio | p |
| --- | --- | --- | --- | --- | --- | --- | --- |
| A | 1 | Ax - Lb | 0.449 | 0.309 | 162 | 1.454 | 0.3161 |
| A | 1 | Ax - Ap | 0.804 | 0.309 | 162 | 2.603 | 0.0271 |
| A | 1 | Lb - Ap | 0.355 | 0.309 | 162 | 1.15 | 0.485 |
| B | 1 | Ax - Lb | 0.51 | 0.309 | 162 | 1.65 | 0.2278 |
| B | 1 | Ax - Ap | 1.398 | 0.309 | 162 | 4.525 | <.0001 |
| B | 1 | Lb - Ap | 0.888 | 0.309 | 162 | 2.875 | 0.0127 |
| A | 2 | Ax - Lb | -0.405 | 0.309 | 162 | -1.311 | 0.391 |
| A | 2 | Ax - Ap | 3.686 | 0.309 | 162 | 11.93 | <.0001 |
| A | 2 | Lb - Ap | 4.091 | 0.309 | 162 | 13.241 | <.0001 |
| B | 2 | Ax - Lb | 0.443 | 0.309 | 162 | 1.434 | 0.326 |
| B | 2 | Ax - Ap | 3.078 | 0.309 | 162 | 9.961 | <.0001 |
| B | 2 | Lb - Ap | 2.635 | 0.309 | 162 | 8.528 | <.0001 |
| A | None | Ax - Lb | 0.925 | 0.309 | 162 | 2.995 | 0.0089 |
| A | None | Ax - Ap | 1.328 | 0.309 | 162 | 4.298 | 0.0001 |
| A | None | Lb - Ap | 0.403 | 0.309 | 162 | 1.303 | 0.3956 |
| B | None | Ax - Lb | -0.266 | 0.309 | 162 | -0.862 | 0.6649 |
| B | None | Ax - Ap | 0.666 | 0.309 | 162 | 2.156 | 0.082 |
| B | None | Lb - Ap | 0.933 | 0.309 | 162 | 3.018 | 0.0083 |

<sup>†</sup> Ax = axenic, Ap = *Acetobacter pomorum*, Lb = *Laevilactobacillus brevis*

## Conclusions

Our study suggests that microbial regulation of fly TAG is highly dependent not only on media preservatives or constituent yeast batch, but also the yeast*preservative interaction. A specific combination of yeast and preservative formula was even sufficient to reverse the effect of microbial elimination, producing a distinct experimental outcome. Preservative formula interfered with microbial effects particularly strongly, with potential to block microbial regulation of host TAG. These overlooked factors appear to be significant determinants of microbiota-dependent fly phenotypes, as well as major causes of microbiota-independent variation. Factors that we have not measured, such as dietary sugar (15) may further influence these complex interactions.

Our results have implications for future fly research. Sparse methodological detailing of diet is a persistent problem, e.g. with methods reporting “standard media”, when media can in fact vary widely among labs. Preservatives are sometimes not reported, and yeast batch variation receives little attention in the lab or literature. Yet our results indicate that these variables can determine experimental outcomes, with implications for repeatability. Our results are consistent with the suggestion that variability among labs may result from yeast batch variation (23). We suggest that diet standardization (e.g. chemically-defined diet) may mitigate these potential confounding factors. Further studies are required to systematically determine how experimental contexts determine outcomes of manipulating the microbiota.

## Acknowledgments

This work was funded by BBSRC (BB/W510658/1), a UKRI Future Leaders Fellowship (MR/S033939/1), and a Lord Kelvin Adam Smith Fellowship from the University of Glasgow. DRS and AJD designed the research, DRS performed the research, DRS and AJD analyzed the data, DRS drafted the manuscript, and AJD edited the manuscript. We thank Nathan Woodling for helpful comments on the manuscript.

